## Supplementary material for "Asymmetric Histone Inheritance Regulates Olfactory Stem Cell Fates During Regeneration": Ma_Supplementary information: Figure S1-S9

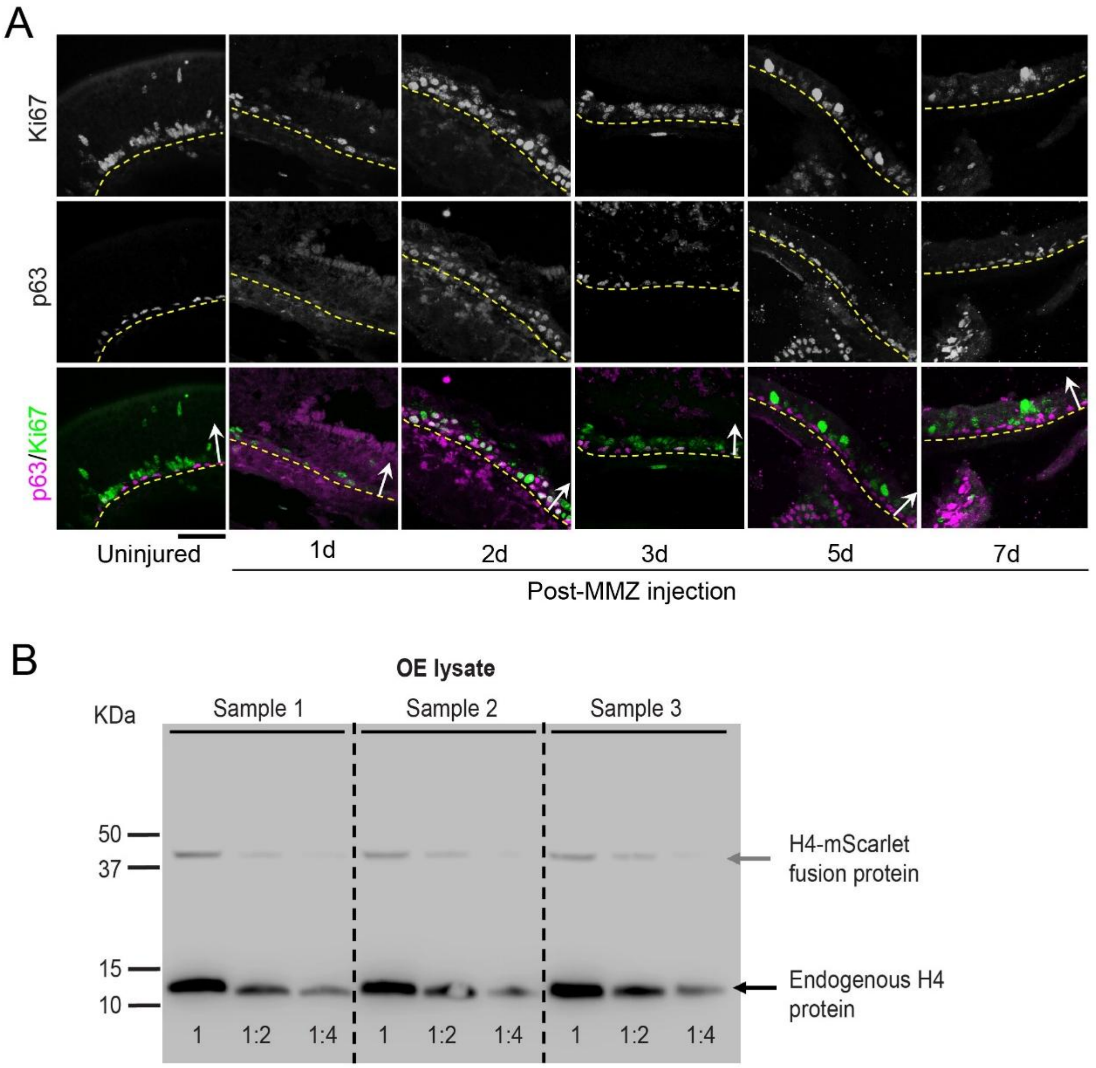


**Supplementary Fig. S1. Post-injury proliferation dynamics of p63+ HBCs and immunoblotting analysis of tagged histone H4.** (**A**) Dynamics of Ki67 labeling in p63+ HBCs during early OE regeneration post-MMZ injection. Tissue sections were from uninjured OE (left panel) and injured OE at Days 1, 2, 3, 5 and 7 post-MMZ injection (right panels). The dotted lines indicate the basal membrane. The arrows point toward the epithelium surface. Related to Fig. 1**B**. Scale bar: 50 µm. (**B**) Immunoblot of H4mS;rtTA OE lysate on Day 2 post-MMZ injection with anti-H4 antibody from three different biological duplicates. The three lanes represent the original OE lysate, 1:2 and 1:4 dilution of the original lysate, respectively. Top band, H4-mScarlet fusion protein, ~40 KD; Bottom band, endogenous H4 protein, ~11 KD.

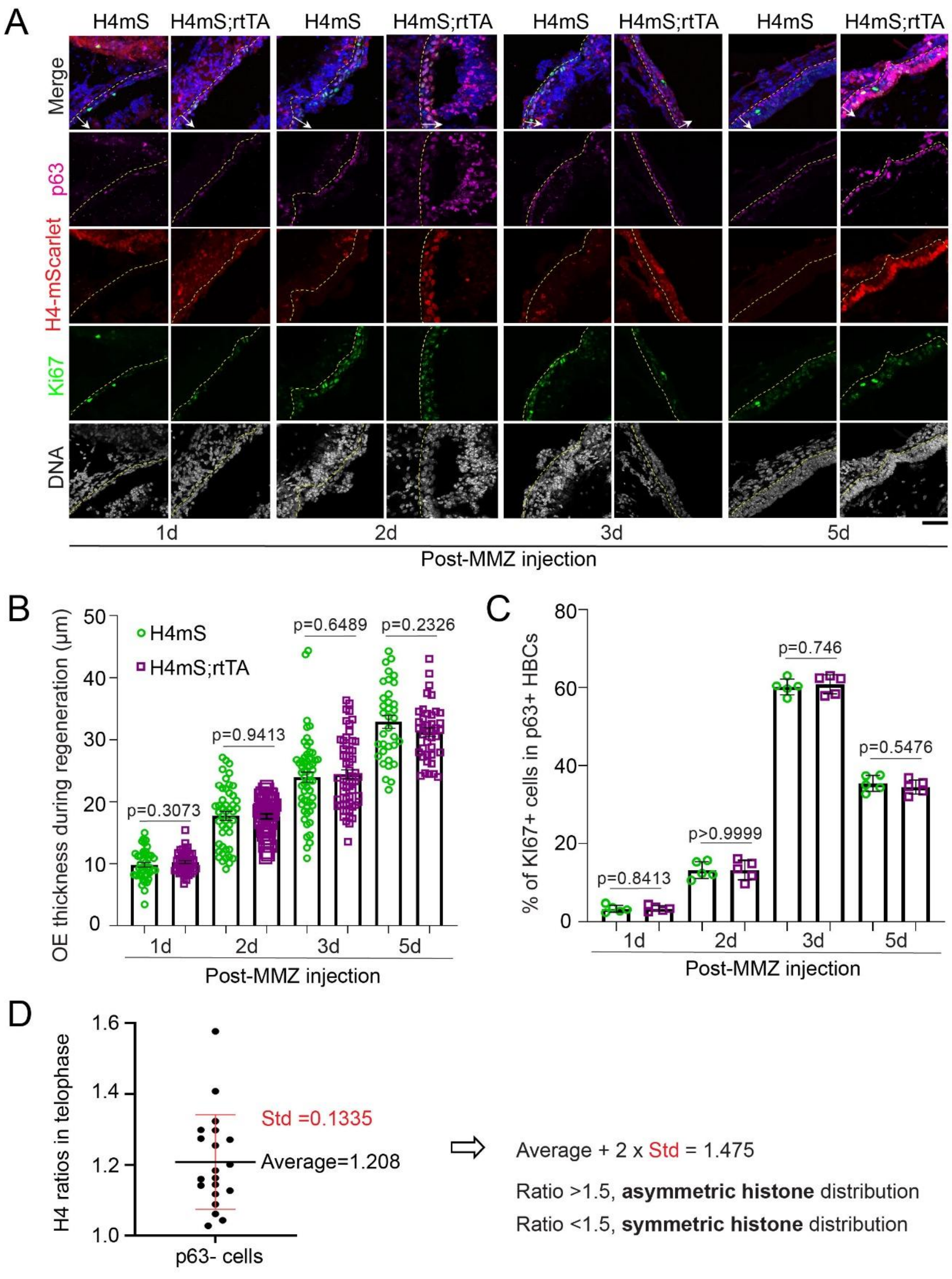
A
H4mS
H4mS;rtTA
H4mS
H4mS;rtTA
H4mS
H4mS;rtTA
H4mS
H4mS;rtTA
Merge
p63
H4-mScarlet
Ki67
DNA
1d
2d
3d
5d
Post-MMZ injection
B
H4mS
H4mS;rtTA
OE thickness during regeneration (μm)
50
40
30
20
10
0
p=0.3073
p=0.9413
p=0.6489
p=0.2326
1d
2d
3d
5d
Post-MMZ injection
C
% of KI67+ cells in p63+ HBCs
80
60
40
20
0
p=0.8413
p>0.9999
p=0.746
p=0.5476
1d
2d
3d
5d
Post-MMZ injection
D
H4 ratios in telophase
1.6
1.4
1.2
1.0
Std =0.1335
Average=1.208
p63- cells
Average + 2 x Std = 1.475
Ratio >1.5, asymmetric histone distribution
Ratio <1.5, symmetric histone distribution

**Supplementary Fig. S2. H4mS;rtTA mice show normal HBCs activation during early OE regeneration.** (**A**) Dynamics of HBC activation and proliferation at Days 1, 2, 3 and 5 post-MMZ injection between H4mS control (H4mS without rtTA) and H4mS;rtTA mice. The dotted line indicates the basal membrane. The arrow points toward the epithelium surface. Scale bar: 50 μm. (**B**) Percentages of Ki67+ cells in p63+ HBCs. Statistical differences according to two-sided Mann-Whitney test. $n = 3$ biological replicates and error bars represent mean ± SEM. (**C**) Quantification of OE thickness. Statistical differences according to two-sided Mann-Whitney test. (**D**) Determination of the cutoff for Histone H4 asymmetry. Left panel shows the ratio of H4 levels between two sister chromatids from p63-negative (p63-) telophase cells from OE sections of H4mS;rtTA mice at Day 2 post-MMZ injection. With the average + 2 × Std (1.475) as the reference, 1.5-fold difference was chosen as the cutoff for assigning asymmetric histone distribution. This cutoff is a confident call and consistent with published cutoff for calling protein asymmetry[61]. Related to Fig. 1H-J, 2C'-F', 3D'-F', 4L, 5C, E, J, 6C, S4D-E, S6C-D and S7I.

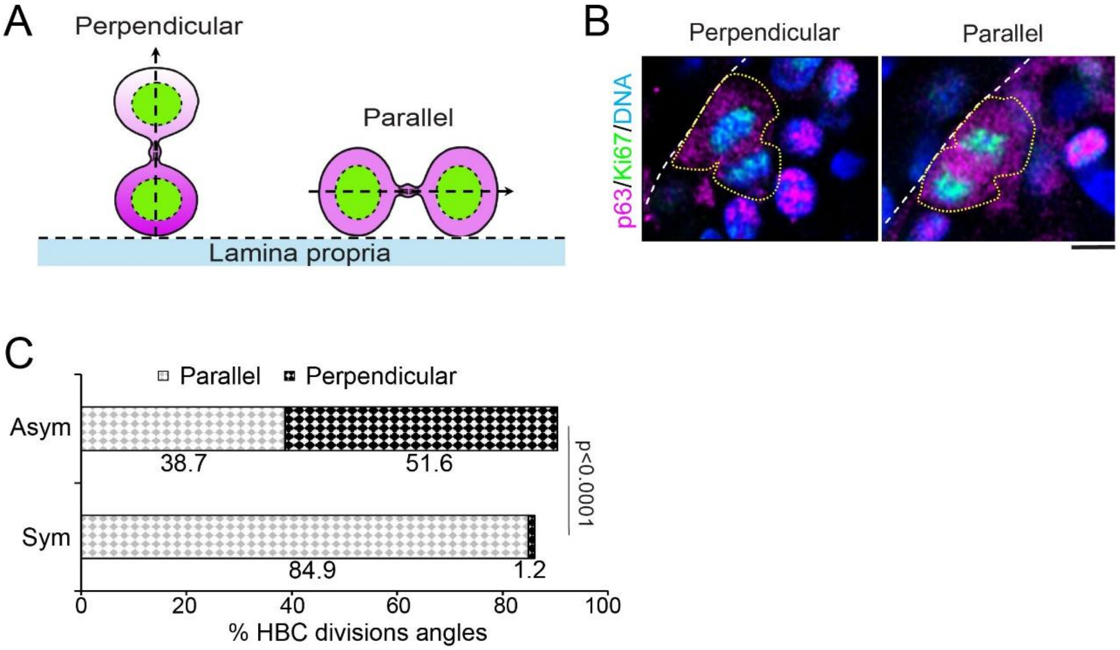


**Supplementary Fig. S3. Division angles of p63+ HBCs during early OE regeneration.** (**A**) Illustration of perpendicular and parallel divisions. The division angle is the angle between the division axis (dashed arrows) and the basal membrane. 0-30°: parallel division; 60-90°: perpendicular division. (**B**) Images of a perpendicular HBC division (left panel) and a parallel HBC division (right panel) with staining of p63, Ki67 and DNA. Scale bar: 5 μm. (**C**) Percentages of perpendicular and parallel divisions in p63+ anaphase/telophase HBCs that show asymmetric or symmetric p63 distribution ($N_{Asym}$ = 31, $N_{Sym}$ = 86) on Day 2 post-MMZ injection. $n = 3$ biological replicates and statistical differences according to Chi-square test.

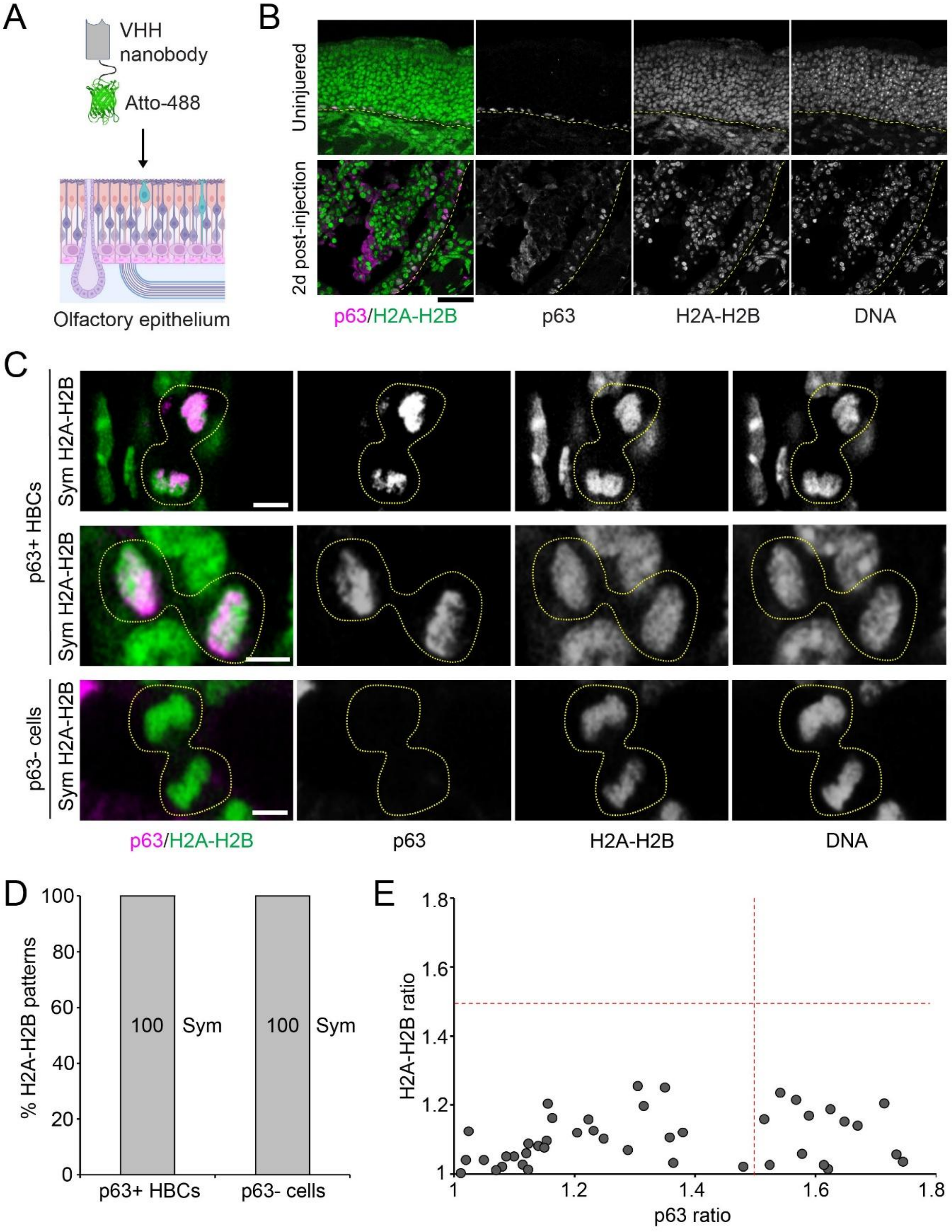
A
VHH nanobody
Atto-488
Olfactory epithelium
B
Uninjuered
2d post-injection
p63/H2A-H2B
p63
H2A-H2B
DNA
C
p63+ HBCs
Sym H2A-H2B
Sym H2A-H2B
p63- cells
Sym H2A-H2B
p63/H2A-H2B
p63
H2A-H2B
DNA
D
% H2A-H2B patterns
100
80
60
40
20
0
100
Sym
100
Sym
p63+ HBCs
p63- cells
E
H2A-H2B ratio
1.8
1.6
1.4
1.2
1
1
1.2
1.4
1.6
1.8
p63 ratio

**Supplementary Fig. S4. Symmetric distribution of histone H2A-H2B in dividing HBCs.** (**A**) Illustration of H2A-H2B nanobody staining in OE sections. (**B**) Staining of H2A-H2B and p63 in uninjured and injured OE (Day 2 post-MMZ injection). The dotted lines indicate the basal membrane. The arrows point toward the epithelium surface. (**C**) H2A-H2B distribution patterns in telophase HBCs. (top) a p63+ HBC with symmetric H2A-H2B distribution and asymmetric p63 distribution, (middle) a p63+ HBC with symmetric H2A-H2B distribution and symmetric p63 distribution, (bottom) a p63-negative (p63-) HBC with symmetric H2A-H2B distribution. (**D**) Ratios of H2A-H2B distribution patterns in p63+ HBCs (N = 45) and p63- cells (N = 15) at telophase. Cutoff of asymmetry = 1.5. (**E**) Correlation of H2A-H2B ratios and p63 ratios between two sister chromatids in p63+ telophase HBC. R=0.174 (N = 45). Scale bars: 50 µm in (**B**); 5 µm in (**C**). The cartoons are created by BioRender.com.

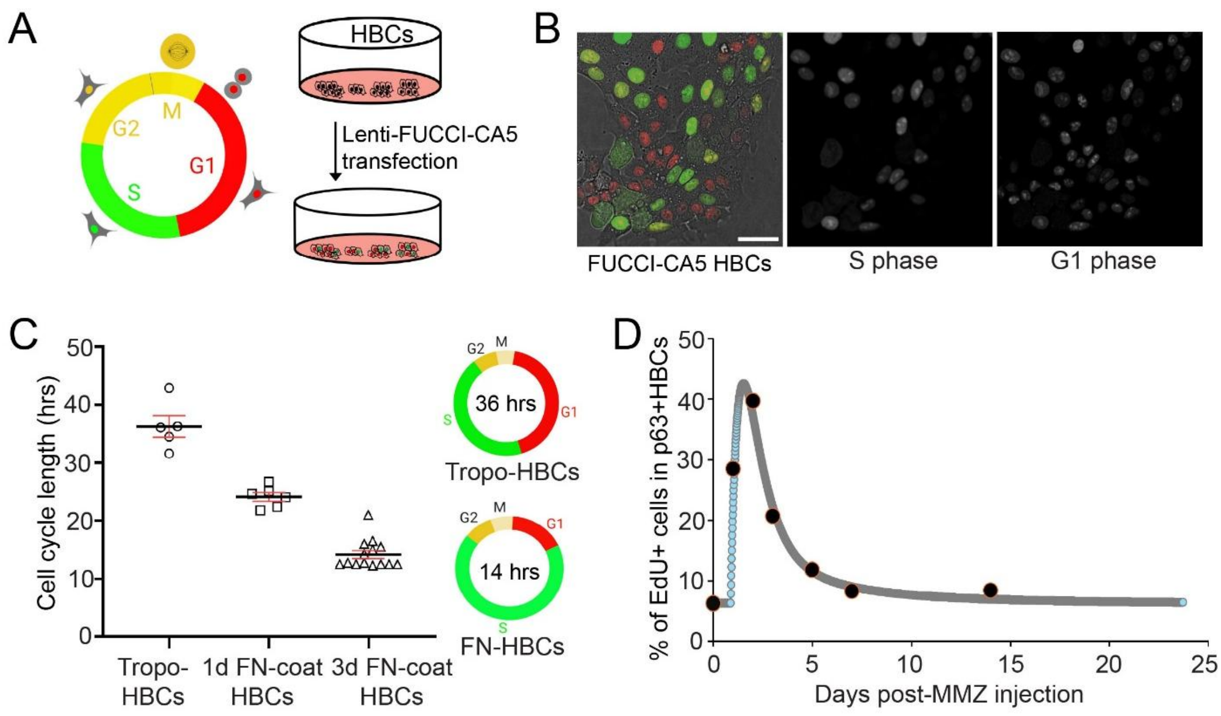


**Supplementary Fig. S5. Primary cultured HBCs recapitulate the *in vivo* HBC activation and proliferation.** (**A**) Illustration of FUCCI-CA5 reporter system and transfection of FUCCI-CA5 plasmid into primary cultured HBCs to track cell cycle progression with live imaging. (**B**) Images of FUCCI-CA5 transfected HBCs with G1 phase cells (red in the merged panel) and S phase cells (green in the merged panel). (**C**) Cell cycle lengths of HBCs from Tropoelastin (Tropo) and Fibronectin (FN) coated conditions. (**D**) Mathematical modeling of cell cycle phases from primary cultured HBCs (curve) fits the *in vivo* HBC proliferation dynamics during early OE regeneration measured by EdU labeling (dots). Scale bars: 25 μm in (**B**). The cartoon elements are created by BioRender.com.

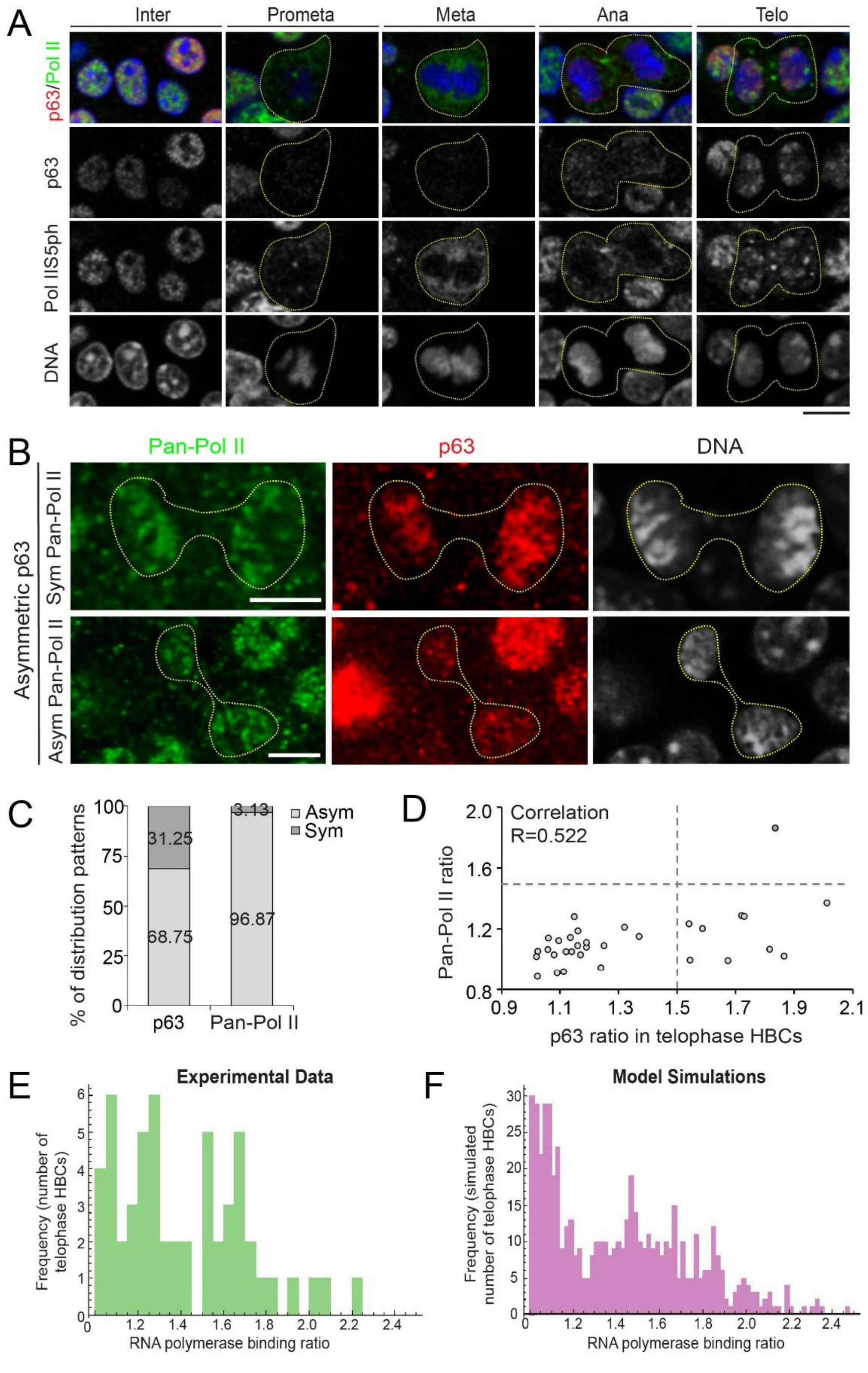
A
Inter
Prometa
Meta
Ana
Telo
p63/Pol II
p63
Pol IIS5ph
DNA
B
Pan-Pol II
p63
DNA
Asymmetric p63
Sym Pan-Pol II
Asym Pan-Pol II
C
% of distribution patterns
Asym
Sym
31.25
68.75
3.13
96.87
p63
Pan-Pol II
D
Correlation
R=0.522
Pan-Pol II ratio
p63 ratio in telophase HBCs
E
Experimental Data
Frequency (number of telophase HBCs)
RNA polymerase binding ratio
F
Model Simulations
Frequency (simulated number of telophase HBCs)
RNA polymerase binding ratio

**Supplementary Fig. S6. RNA Pol II distribution and mathematical modeling of differential transcription re-initiation.** (**A**) Dynamics of RNA Pol IIS5ph distribution during interphase and mitosis. (**B**) Immunostaining of unphosphorylated Pol II (Pan-Pol II) and p63 in telophase HBCs. Top panel, symmetric distribution pattern of Pan-Pol II in a telophase HBC with asymmetric p63 distribution. Bottom panel, asymmetric distribution pattern of Pan-Pol II in a telophase HBC with asymmetric p63 distribution. (**C**) Quantification of the ratio of unphosphorylated Pol II and p63 in telophase HBCs (N = 32). The cutoff for unphosphorylated Pol II and p63 asymmetry is 1.5. (**D**) The correlation plot of ratios of unphosphorylated Pol II and p63 in each telophase HBC. R = 0.522. (**E** and **F**) Mathematical modeling of differential transcription re-initiation elutes differential binding affinity of RNA Pol II in telophase HBCs. (**E**) Experimental data of RNA Pol IIS2ph distribution patterns from telophase HBCs based on the immunostaining results. Frequency: number of telophase HBCs. (**F**) Histogram of mathematical simulation of differential transcription re-initiation with experimental data of RNA Pol IIS2ph distribution data. The modeling indicates a 60% difference in RNA Pol II transcription binding affinity between two sister chromatids of telophase HBCs. Frequency: simulated number of telophase HBCs. Scale bars: 10 μm in (**A**); 5 μm in (**B**).

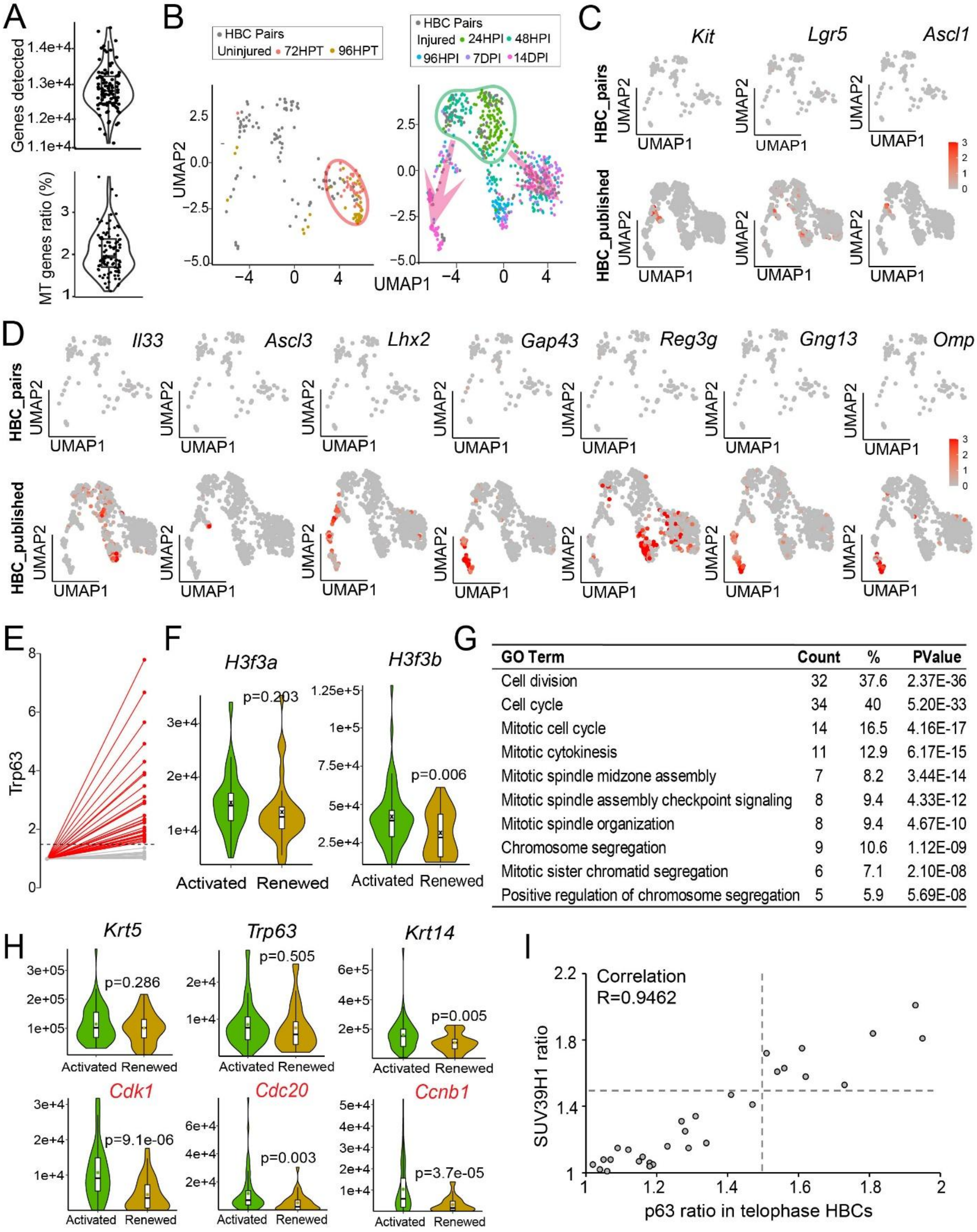


| GO Term | Count | % | PValue |
|---|---|---|---|
| Cell division | 32 | 37.6 | 2.37E-36 |
| Cell cycle | 34 | 40 | 5.20E-33 |
| Mitotic cell cycle | 14 | 16.5 | 4.16E-17 |
| Mitotic cytokinesis | 11 | 12.9 | 6.17E-15 |
| Mitotic spindle midzone assembly | 7 | 8.2 | 3.44E-14 |
| Mitotic spindle assembly checkpoint signaling | 8 | 9.4 | 4.33E-12 |
| Mitotic spindle organization | 8 | 9.4 | 4.67E-10 |
| Chromosome segregation | 9 | 10.6 | 1.12E-09 |
| Mitotic sister chromatid segregation | 6 | 7.1 | 2.10E-08 |
| Positive regulation of chromosome segregation | 5 | 5.9 | 5.69E-08 |

**Supplementary Fig. S7. scRNA-seq identifies gene expression patterns of paired HBC daughters.** (**A**) Combined violin and box plots showing the total number of genes detected (up panel) and the mitochondrial (MT) genes ratio (bottom panel) in the scRNA-seq analysis. Each dot represents data from a single cell (N = 96). (**B**) Left panel, comparison of paired-daughter HBC scRNA-seq data with published HBC scRNA-seq data from uninjured OE (GSE99251). Notice that most HBCs from uninjured OE are resting HBCs (circled in red). Right panel, comparison of paired-daughter HBC scRNA-seq data with published scRNA-seq data from injured OE (GSE95601) through regeneration. The cultured HBCs were collected from injured OE at Day 2 post-MMZ injection and cultured first with Tropoelastin coating and then Fibronectin coating. Notice that most cultured HBCs (circled in green) overlap with 24-hour and 48-hour post injury groups of HBCs from the *in vivo* samples. Arrow indicates multilineage fate priming. (**C**) Expression of GBCs marker genes *Ascl1*, *Kit* and *Lgr5* in HBC pairs (top panels) and published datasets (lower panels). (**D**) Expression of marker genes from different OE cell types, including sustentacular cell marker *Il33*; microvillous cell marker *Ascl3*; immediate neuronal precursor marker *Lhx2*; immature olfactory sensor neuron markers *Gap43*, *Reg3g* and mature olfactory sensor neuron markers *Gng13*, *Omp* in HBC pairs (top panels) and published datasets (lower panels). (**E**) Expression of P63 transcripts (*Trp63*) in individual pair. Red lines indicate asymmetric pairs (N = 35). Grey lines indicate symmetric pairs (N = 13). The asymmetric cutoff for *Trp63* is 1.5. (**F**) Expression of *H3f3a* and *H3f3b* in activated and renewed HBC cells. (**G**) Gene ontology analysis of paired-daughter HBC scRNA-seq data for biological processes related to genes highly expressed in activated HBCs. (**H**) Expression of *Krt5*, *Trp63*, *Krt14* and *Cdk1*, *Cdc20*, *Ccnb1* (labeled in red, data from Fig. 4J) in combined activated and renewed HBCs from the paired-daughter HBC dataset. In (**F**) and (**H**), all box plots include the median and data between the 25th and 75th percentile, with the whiskers indicating lower and upper extremes after removing outliers. The black 'x' within each box shows the average within each group. Two-tailed t-test was used to calculate P-values. (**I**) Correlation of SUV39H1 ratios and p63 ratios between two sister chromatids in p63+ telophase HBC. The cutoff for p63 and SUV39H1 asymmetry is 1.5. R = 0.9462 (N = 29).

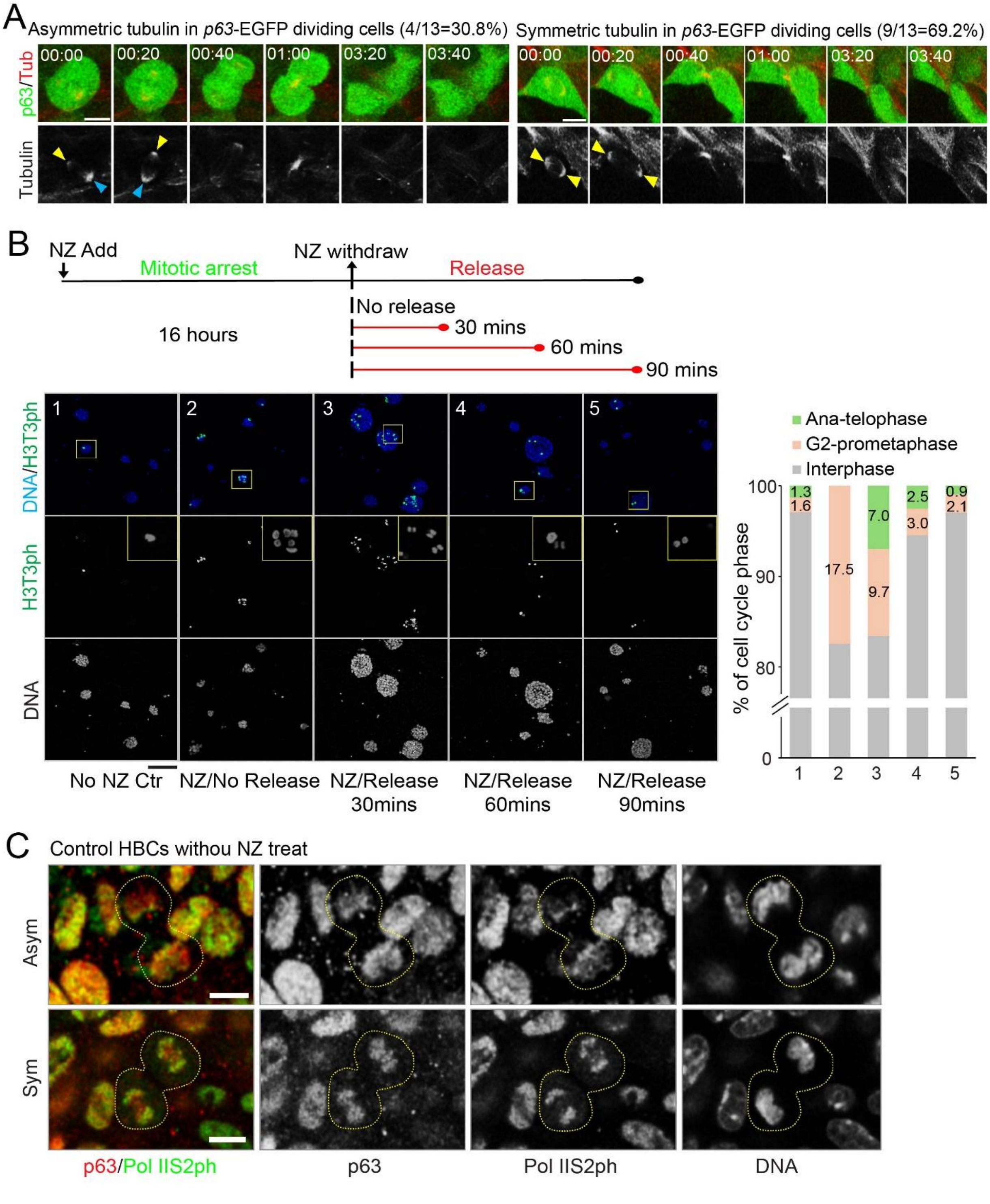
A
Asymmetric tubulin in p63-EGFP dividing cells (4/13=30.8%)
Symmetric tubulin in p63-EGFP dividing cells (9/13=69.2%)
p63/Tub
Tubulin
00:00
00:20
00:40
01:00
03:20
03:40
B
NZ Add
Mitotic arrest
NZ withdraw
Release
No release
16 hours
30 mins
60 mins
90 mins
DNA/H3T3ph
H3T3ph
DNA
No NZ Ctr
NZ/No Release
NZ/Release 30mins
NZ/Release 60mins
NZ/Release 90mins
Ana-telophase
G2-prometaphase
Interphase
% of cell cycle phase
1.3
1.6
17.5
7.0
9.7
2.5
3.0
0.9
2.1
C
Control HBCs withou NZ treat
Asym
Sym
p63/Pol IIS2ph
p63
Pol IIS2ph
DNA

**Supplementary Fig. S8. Asymmetric microtubule and RNA Pol II distribution in dividing HBCs.** (**A**) Live cell imaging of microtubule activity using the live tubulin dye SiR-Tubulin (red) and *p63*-EGFP transcriptional reporter (green) during HBCs divisions (N=13). Image interval, 20 minutes. (Left panel) Example of a dividing HBC showing asymmetric microtubule activity and differential level of p63-EGFP. 4 of 13 dividing HBCs (30.8%) exhibited asymmetric microtubule activity. Blue arrows indicate higher microtubule activity; yellow arrows indicate lower microtubule activity. (Right panel) Example of a dividing HBC showing symmetric microtubule activity and comparable level of p63-EGFP. 9 of 13 dividing HBCs (69.2%) exhibited symmetric microtubule activity. Yellow arrows indicate equal microtubule activities. (**B**) Assessment of mitotic progression after Nocodazole (NZ) treatment with different releasing time in primary cultured HBCs. HBCs were cultured with NZ for 16 hours followed by release and imaging. Before NZ treatment, ~1% of HBCs were in anaphase/telophase. After 16-hour treatment, ~18% of HBCs were arrested at prometaphase. Upon release, HBCs continued mitotic progression. At 30-minute releasing time, ~7% of HBCs were found in anaphase/telophase (right panel). At 60- and 90-minute releasing time, ~3% and ~1% of HBCs were found in anaphase/telophase, respectively (right panel). The mitotic HBCs are indicated by H3T3ph and DNA. (**C**) Asymmetrically (top panel) and symmetrically (bottom panel) dividing HBCs at telophase in control group without NZ treatment. Scale bars: 5 µm in (**A**); 100 µm in (**B**); 5 µm in (**C**).

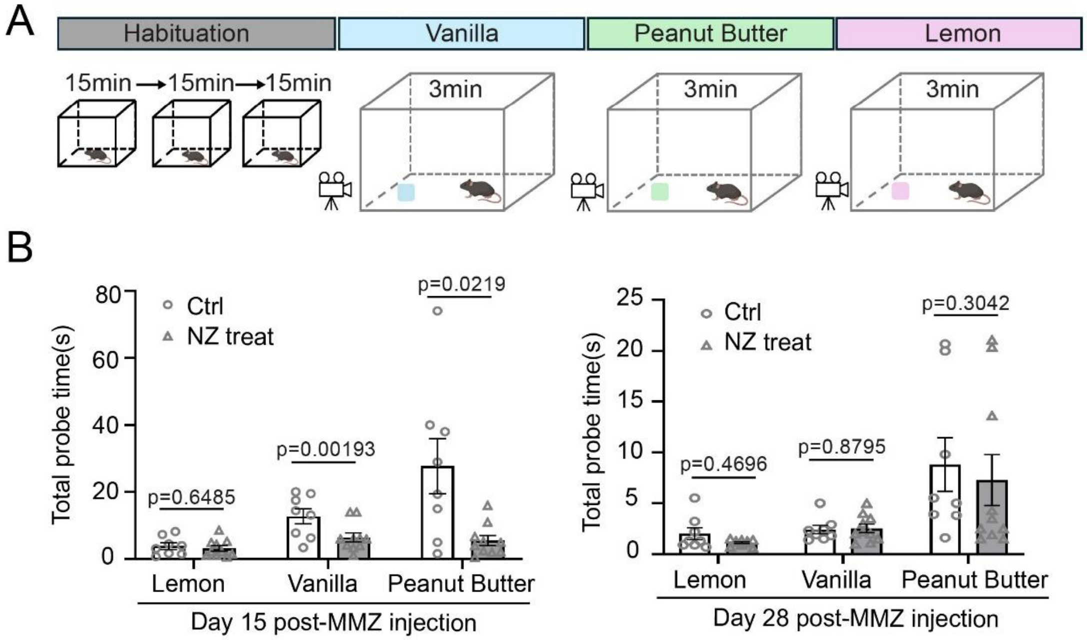


**Supplementary Fig. S9. Olfactory preference test in NZ-treated mice**. (**A**) Illustration of olfactory preference test in NZ-treated mice and non-NZ treated control mice. The mice were tested as a cohort, with each mouse exposed to an odor source for 3 minutes. The total probe time for each mouse was quantified. The cohort then proceeded to the next odor only after all mice in the cohort had completed the test for the current odor. $N_{Ctrl}$ = 8 mice; $N_{NZ\ treat}$ = 10 mice. (**B**) Quantification of total probe time from the olfactory preference test on Day 15 and Day 28 post-MMZ injection with NZ treatment. Total probe time in Ctrl mice on Day 15: Lemon = 3.88 ± 0.96, Vanilla = 12.75 ± 2.22, Peanut butter = 27.74 ± 8.27; Total probe time in NZ-treated mice on Day 15: Lemon = 3.19 ± 0.81, Vanilla = 6.48 ± 1.35, Peanut butter = 5.51 ± 1.50. Total probe time in Ctrl mice on Day 28: Lemon = 2.00 ± 0.58, Vanilla = 2.40 ± 0.41, Peanut butter = 8.79 ± 2.65; Total probe time in NZ-treated mice on Day 28: Lemon = 1.14 ± 0.10, Vanilla = 2.53 ± 0.42, Peanut butter = 7.28 ± 2.51. Values represent average ± SEM. Statistical differences according to two-sided Mann-Whitney test. The cartoons are created by PowerPoint.
